# Evidence for a compact σ^70^ conformation *in vitro* and *in vivo*

**DOI:** 10.1101/2022.10.14.512049

**Authors:** Khalil Joron, Joanna Zamel, Nir Kalisman, Eitan Lerner

## Abstract

The initiation of transcription in Escherichia coli (*E. coli*) is facilitated by promoter specificity factors, also known as σ factors, which may bind a promoter only as part of a complex with RNA polymerase (RNAP). By performing *in vitro* cross-linking mass spectrometry (CL-MS) of apo-σ^70^, we reveal structural features suggesting a compact conformation compared to the known RNAP-bound extended conformation. Then, we validate the existence of the compact conformation using *in vivo* CL-MS by identifying cross-links similar to those found *in vitro*, which deviate from the extended conformation only during the stationary phase of bacterial growth. Conclusively, we provide information in support of a compact conformation of apo-σ^70^ that exists in live cells, which might represent a transcriptionally-inactive form that can be activated upon binding to RNAP.

## Introduction

σ promoter specificity factors facilitate the specific binding of RNAP to gene promoters in bacteria^1,2^. Among the different types of σ factors in *E. coli*, σ^70^ binds promoters of house-keeping genes under optimal conditions at the logarithmic phase of bacterial growth^3^. The mechanism of initiating DNA transcription involves the binding of σ^70^ to dsDNA promoter only when σ^70^ is bound to the core RNAP complex^4^. In the RNAP complex, σ^70^ is found in a unique structural organization, in which σ^70^ regions 2 and 4 (σR2 and σR4, respectively) are distant from each other^4^, to expose its DNA-binding residues, and by that facilitate transcription initiation^5^. The formation of the RNAP-promoter closed (RP_C_) complex is followed by a cascade of DNA isomerization events^6–10^, which end with a stretch of 10-12 melted bases of promoter DNA upstream to the transcription start site, which forms the DNA transcription bubble, that is later stabilized in the RNAP-promoter open (RP_O_) complex^11,12^. Importantly, throughout transcription initiation σ^70^ remains part of the holoenzyme complex and retains the extended conformation. Nonetheless, as all parts of the transcription initiation complex, including σ^70^, are translated and formed separately, σ^70^ may exist in bacteria also in an unbound apo form, at least as long as it is not bound to RNAP. Apo-σ^70^ can also bind anti-σ factors, which repress transcription by competing with RNAP on the interaction with σ^70^, and even by sequestering σ^70^ out of the cytoplasm^13–15^. However, different factors may lead to the release of σ^70^ from its interaction with anti-σ factors, and back into the cytoplasm, until it rebinds RNAP^16–19^. Therefore, this process, among others, might dictate the lifetime in which σ^70^ exists in an unbound apo form.

Structurally, the protein data bank (PDB) includes many entries of high-resolution structures of transcription initiation complexes, where σ^70^ is present as a subunit^4,8,20–30^. Exploring the PDB for apo-σ^70^ identifies only a few *E. coli*-related structures of its separate regions, such as σ^70^ region 4 (i.e., σR4) bound to several transcription factors^31–33^, the unbound σ^70^ region 2 (i.e., σR2)^34,35^, as well as structures of regions in house-keeping σ factors from bacteria other than *E. coli*^35,36^. However, a structural description of fulllength apo-σ^70^ has not yet been reported. Nevertheless, previous *in vitro* biochemical works showed that apo-σ^70^ adopts a predominant distinct compact conformation^37–40^, which can undergo an overall structural reorganization upon binding core RNAP. Binding induces a conformational change from an unbound compact state to a RNAP-bound extended state^41–45^.

In this work, we provide evidence for the existence of the compact apo-σ^70^ conformation both *in vitro* and *in vivo*. First, we perform *in vitro* single-molecule Förster resonance energy transfer (smFRET)^46,47^ experiments to verify that apo-σ^70^ is found predominantly in a compact conformation. Moreover, we also find that the presence of a promoter DNA alone (i.e., in the absence of RNAP) can induce only a minor population shift towards the extended conformation of σ^70^. Relying on smFRET data, we further study the structural features of apo-σ^70^ by performing *in vitro* cross-linking mass-spectrometry (CL-MS) ^48–53^. Comparison of the resolved residue proximities against the PDB full-length σ^70^ structures confirms the description of apo-σ^70^ as having a compact conformation. Furthermore, we show a global decrease in the distance between σR2 and σR4, which could support a DNA-binding auto-inhibitory mechanism, for preventing DNA binding not in the context of transcription.

Finally, results from *in vivo* CL-MS experiments on σ^70^ in *E. coli*, report the abundance of inter-residue proximities that cannot be explained by the extended σ^70^ PDB structures. Interestingly, the compact apo-σ^70^ conformation exists not in the logarithmic bacterial growth phase when σ^70^ is recruited for transcription, but rather in the stationary phase when alternative σ factors are recruited for transcription to replace σ^70^.

### *In vitro* single-molecule FRET of apo-σ^70^ reveals a predominant compact conformation

To study the inter-region conformational dynamics of σ^70^, we doubly labeled σR2 and σR4 with fluorescent dyes for smFRET measurements, as shown before^37^(see Methods). Importantly, using an smFRET hybridizing probe-based transcription assay, presented previously by Weiss and co-workers^54,55^, we show that both the *wt* and the mutant used for dye-labeling are transcriptionally-active (Fig. S1). The double-labeled σ^70^ has an expected FRET efficiency of ∼0.5 when it is a subunit of the RNAP holoenzyme structure<u>^5^.</u> SmFRET measurements result in a predominant FRET sub-population with higher FRET efficiency (Fig. 1, a-c), which point towards a conformation of σ^70^ where regions 2 and 4 are in close proximity, consistent with previously published data of similar smFRET measurements<u>^5^.</u> Further, we report the distribution of raw FRET values (E_raw_) of single σ^70^ molecules, probed for freely-diffusing molecules through a confocal spot within few ms, in which they can undergo conformational changes at the diffusion timescale or faster. To report the underlying conformational states and the potential within-burst conformational dynamics, we analyze the smFRET burst data using multi-parameter photon-by-photon hidden Markov modeling (mpH^2^MM)^56^ (Fig. 1, d-f). Apo-σ^70^ transitions between four subpopulations, with one being an acceptor-photoblinked sub-population, and hence does not report FRET-relevant information (Fig. 1, d, purple dots), and three more sub-populations with both donor and acceptor fluorescently-active (Fig. 1, d, blue, red and green dots). The rates of the transitions between the three FRET sub-populations are in the ms timescale, where only rates of 20 s^-1^ or higher can be considered as arising from within-burst dynamics recorded in the experiment, due to burst durations of 50 ms at most (Table S1). Two out of the three FRET sub-populations exhibit almost equal mean E_raw_ values (Fig. 1, d, black dots within the group of blue and red dots) and together represent one predominant FRET sub-population (Fig. 1, d, blue and red dots). The third minorly-populated FRET sub-population (Fig. 1, d, green dots) exhibits a slightly lower mean E_raw_ (Fig. 1, d, black dot within the group of green dots). These results suggest a predominant compact σ^70^ conformation and a minorly-populated slightly-less compact σ^70^ conformation. Incubating the dye-labeled apo-σ^70^ in the presence of 2 μM dsDNA *lacCONS*^54,57^ promoter (Fig. 1, c, f), gives rise to a small fraction of σ^70^ molecules in a FRET sub-population (Fig. 1, f, green dots) at lower mean E_raw_ (Fig. 1, f, black dot within the group of green dots) compared to the same minorly-populated sub-population in apo-σ^70^. These results correspond to a further increase in the distance between σR2 and σR4 for a low fraction of σ^70^ molecules, whereas the majority of apo-σ^70^ stay in the high FRET sub-population. In the presence of lower concentration of dsDNA *lacCONS* promoter (100 nM; Fig. 1, b, e), this minorly-populated sub-population decreases to values similar to those in the absence of dsDNA (as seen relative to Fig. 1, a). Notably, the dsDNA *lacCONS* promoter should bind σ^70^, when it is a subunit of the RNAP holoenzyme^57,58^. Indeed, in previous smFRET studies involving σ^70^ a full shift towards lower FRET values occurred in the presence of only a few nM of dsDNA, only when in the presence of RNAP^9,10^. Therefore, a small fraction of apo-σ^70^ molecules can undergo a small conformational change in the presence of μM concentrations of dsDNA. The transition rates from the predominant compact conformation represented by the high FRET sub-populations (blue and red) to the slightly-less compact sub-population (green), are in the tens of ms, and become as slow as hundreds of ms in the presence of dsDNA (Table S1). It is important to remember, however, that transition rates slower than 20 s^-1^ only point towards the inability of mpH^2^MM to resolve the slow conformational dynamics in these cases.

**Fig. 1.**
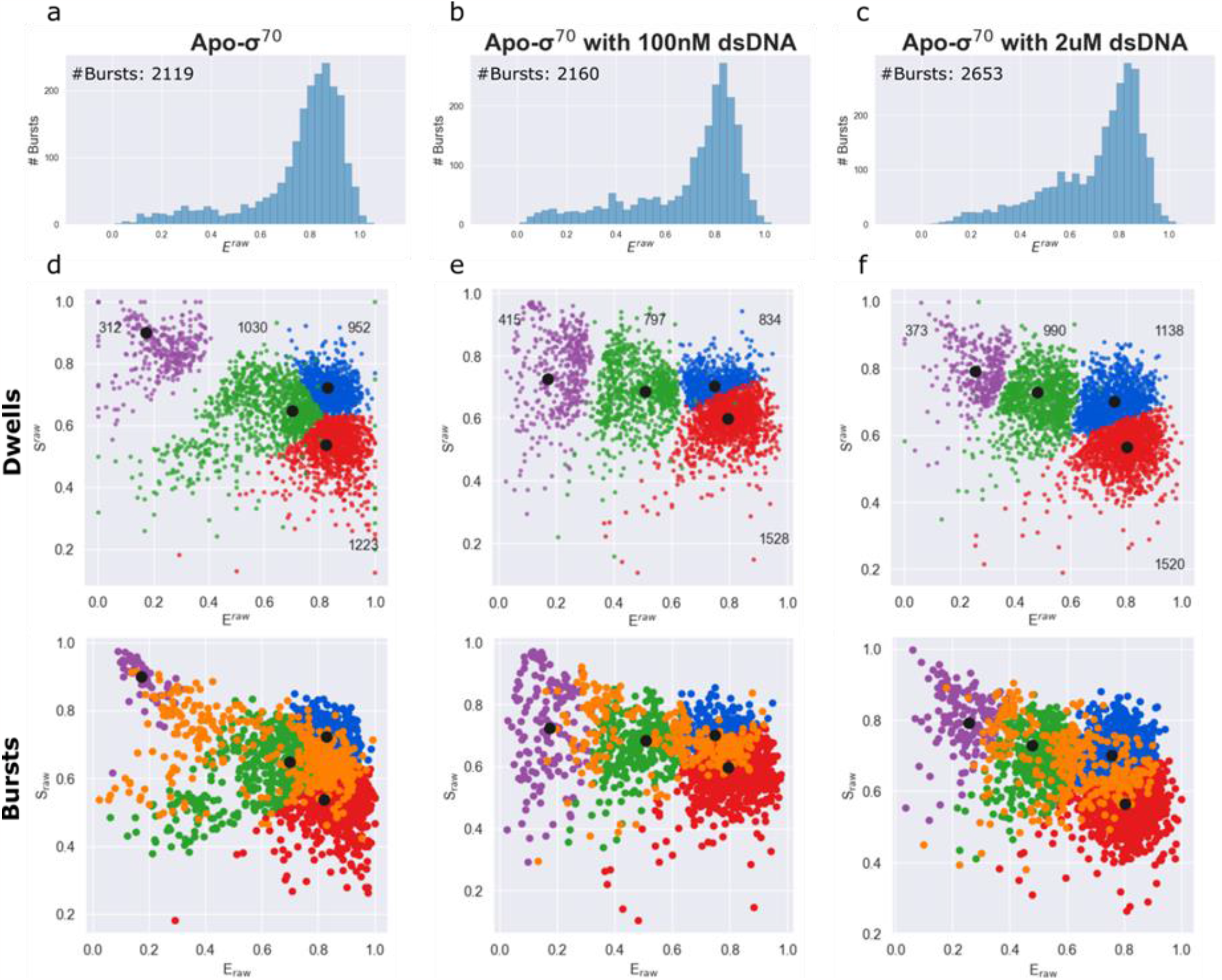
SmFRET analyses of apo-σ^70^ conformations in the absence and presence of promoter dsDNA. (**a-c**) smFRET burst analysis (**d-f**) mpH^2^MM analysis of σ^70^ labeled at residues 442 and 579 with ATTO 550 and ATTO 643, respectively. (**a**) Apo-σ^70^, a major population exists at high raw FRET efficiencies, E_raw_,with a tail towards lower E_raw_ values. (**b**) Apo-σ^70^ with 100 nM dsDNA, exhibits no significant change in the major E_raw_ population. (**c**) Apo-σ^70^ with 2 μM dsDNA, exhibits a slight increase in sub-population with lower E_raw_. (**d-f**) Dwells in states within bursts, (dwell-based analysis; top) and burst-based analysis (bottom), where different sub-populations in different states are shown in blue, red, green and magenta, and the bursts that include transitions between states are shown in orange. (**d**) Apo-σ^70^ – the predominant sub-populations (blue and red) of the compact conformation exhibit dynamics to a minor sub-population (green) of a less compact conformation. (**e**) σ^70^ in the presence of 100 nM dsDNA – the green sub-population is shifted, relative to (d), towards lower mean E_raw_ values suggesting binding to dsDNA influences the conformation of a minor sub-population in σ^70^, while (**f**) in the presence of 2 μM dsDNA shows a higher an increase in the population of the green sub-population, suggesting the dsDNA-bound σ^70^ occurs more often at μM concentrations, yet most of σ^70^ still remains in the high E_raw_ sub-population (in the compact conformation; blue and red). Transition times between the blue or red sub-populations and the green sub-population, and vice versa, are in a few to tens of ms, in the absence or presence of dsDNA. The magenta sub-population can be best described by a pure donor-only sub-population shown using uncorrected raw FRET and ALEX data (E_raw_ and S_raw_).

Overall, the results of the *in vitro* smFRET measurements suggest that apo-σ^70^ is in an equilibrium between two states, a predominant compact conformation and a minorly-populated slightly less compact conformation that is still more compact than the fully-extended-RNAP-bound σ^70^, which is well-described by PDB structures. Therefore, we can assume that C_α_-C_α_ distances between σR2 and σR4 in the predominant conformation of apo-σ^70^ may yield the short proximities that allow cross-linking to occur efficiently. Thus, we set out to describe the structural features of the apo-σ^70^ compact conformation using *in vitro* CL-MS.

### Structural features of the unique compact apo-σ^70^ conformation using *in vitro* CL-MS

We perform *in vitro* cross-linking of purified recombinant *wt* apo-σ^70^ with either a zero-length primary amine to carboxyl coupler, 4-(4,6-dimethoxy-1,3,5-triazin-2-yl)-4-methyl-morpholinium chloride (DMTMM)^50^, or a primary amine to primary amine cross-linker, bis(sulfosuccinimidyl)-substrate (BS^3^)^59^. The cross-linked sample is then digested with trypsin and analyzed using MS (for further information, see Supplementary Information). We attain both short and intermediate scale spatial information, which report the C_α_-C_α_ distances that are <16 or <30 Å for DMTMM and BS^3^, respectively (Table 1). We show that without cross-linking the majority of σ^70^ (∼66%) is in a monomeric form, while the rest exists as higherorder species (Fig. S2, a). As a result of cross-linking, the fraction of monomeric σ^70^ decreases by 6%, without affecting the overall layout of the eluted peaks (Fig. S2, b). Therefore, we suggest that many of the cross-links we achieve originate from the monomeric form. After false detection rate (FDR)-based data filtration (Fig. S3, a & b) according to the fragmentation score of each cross-linked peptide (see Supplementary Information), the highest-ranking cross-links are summarized in supplementary spreadsheet. We test the satisfaction rate of the recovered C_α_-C_α_ distances against the same pairs of residues in existing PDB structures of σ^70^ (Table S2). The interconnectivity between σ^70^ regions (Fig. S4), include pairs of cross-linked residues between σ^70^ regions that are expected to be distant from each other (e.g., σR2 and σR4) and should not generate distances within the distance cutoffs of the cross-linkers used, if based on spatial information given by all PDB structures. Since all existing PDB structures of σ^70^ possess similar features and are almost identical, we display the recovered cross-links using one representative PDB structure of σ^70^ in the RNAP holoenzyme complex (6P1K)^25^ (Fig. 2, a & b), and summarize the results for all PDB structures in a table (Table 1). Out of this comparison, we infer that 43-46% of the BS^3^ and 25-38% of the DMTMM cross-links are satisfied and fit well within the structural coordinates of the pairs of residues. According to PDB structures coordinates most cross-links exceed the C_α_-C_α_ distance range covered by the cross-linkers, suggesting that the PDB structures do not represent all possible monomeric conformations captured in solution during cross-linking. Next, we perform modeling using AlphaFold2^60,61^ and RoseTTAFold^62^, to test whether the expected compact conformation may be predicted via training and based on previously acquired structural data alone. AlphaFold2 was able to predict a more compact conformation than the one present in the PDB (Fig. 3, a-c). Yet, predicted models retain a high violation percentage of the cross-linking data (Table S2, and Fig. 3, d & e), suggesting a more compact conformation of apo-σ^70^ exists and is not available in the structural database, which the algorithms were trained on. The relatively low predicted local distance difference test score of σR3 and σR4 (Fig 3, a), suggest that a more compact conformation which the algorithm was not able to predict may exist. According to the residue-residue alignment confidence plot (Fig. 3, b), the interaction of the σR2/σR3 with σR4 is missing in the AlphaFold2 models, contrary to CL-MS results. Next, we test the satisfaction rate of CL-MS data against the predicted structure models, and find a similar level of satisfaction with up to 36% and up to 50% for the DMTMM and BS^3^ data, respectively (Table S2). Overall, across all PDB structures and predicted structure models of σ^70^, we detect the same cross-linked pairs of residues that cannot be explained by the spatial distances given in these structures. Interestingly, these unique pairs of cross-linked residues include mostly ones connecting σR2/σR3 with σR4 (Table 1), which are the primary DNA-binding regions.

**Table 1.** List of pairs of residues recovered from apo-σ^70^ *in vitro* CL-MS. BS^3^ and DMTMM cross-linked pairs of residues recovered from cross-linking mass spectrometry. Residues are assigned to their originating regions: σ region 1.2 (σR1.2), σ non-essential region (σNER), σ region 2 (σR2), σ region 3 (σR3), and σ region 4 (σR4). Residue numbering is according to the sequence given by the PDB structure 6P1K. Blue/Red/Black - residues that satisfy the pair-of-residue C_α_-C_α_ distance given by the PDB structure, residues that violate the distance and unstructured residues in the PDB structure, respectively.

| BS<sup>3</sup> |  |  | BS<sup>3</sup> |  |  | DMTMM |  |  |
| --- | --- | --- | --- | --- | --- | --- | --- | --- |
| Regions | Residue #1 | Residue #2 | Regions | Residue #1 | Residue #2 | Regions | Residue #1 | Residue #2 |
| σR1.2-σNER | K121 | K257 | σNER-σR4 | K343 | K593 | σR1.2 | T107 | E104 |
| σR.12-σNER | K121 | K264 | σNER-σR4 | K264 | K593 | σR1.2-σR2 | D96 | K376 |
| σR1.2-σNER | K121 | K238 | σNER-σR4 | K236 | K578 | σR1.2-σR2 | E104 | K371 |
| σR1.2-σNER | K121 | S241 | σNER-σR4 | K257 | K593 | σR1.2-σR2 | E104 | S389 |
| σR1.2-σNER | K121 | K220 | σR2 | K371 | K377 | σR1.2-σR2 | E109 | K426 |
| σR1.2-σR2 | K121 | K371 | σR2 | K371 | K376 | σR1.2-σR2 | D125 | K371 |
| σR1.2-σR2 | K121 | K426 | σR2-σR3 | K371 | K462 | σR1.2-σR3 | E109 | K462 |
| σ1.2-σR3 | K121 | K462 | σR2-σR3 | K393 | K462 | σNER | K236 | E335 |
| σR1.2-σR4 | K121 | K578 | σR2-σR3 | K418 | K462 | σNER | K236 | E247 |
| σNER | K220 | K251 | σR2-σR3 | K377 | K462 | σNER | E152 | K257 |
| σNER | K220 | K264 | σR2-σR4 | K377 | S604 | σNER | K296 | D332 |
| σNER | K257 | K264 | σR2-σR4 | K371 | K578 | σNER-σR4 | E343 | D613 |
| σNER | K238 | T244 | σR2-σR4 | K377 | K557 | σNER-σR4 | E247 | K557 |
| σNER | K236 | K264 | σR2-σR4 | K377 | K593 | σNER-σR4 | E349 | K593 |
| σNER | K220 | K257 | σR2-σR4 | K392 | K593 | σR2 | E407 | S442 |
| σNER | K236 | K251 | σR2-σR4 | K377 | K578 | σR2-σR3 | D445 | K462 |
| σNER | K289 | K343 | σR3 | K462 | K496 | σR2-σR4 | K376 | D613 |
| σNER | K238 | S241 | σR3 | K462 | K493 | σR2-σR4 | K392 | D546 |
| σNER | K251 | K264 | σR3-σR4 | K462 | K557 | σR3 | K462 | E491 |
| σNER | Y228 | K236 | σR3-σR4 | K462 | K593 | σR3-σR4 | K462 | D546 |
| σNER | K236 | K257 | σR3-σR4 | K462 | S604 | σR3-σR4 | K493 | D546 |
| σNER | Y228 | K257 | σR3-σR4 | K462 | K578 | σR3-σR4 | K493 | D613 |
| σNER | K220 | S253 | σR3-σR4 | K493 | K557 | σR3-σR4 | K462 | D613 |
| σNER-σR2 | K257 | K371 | σR3-σR4 | K496 | K557 | σR4 | D546 | K557 |
| σNER-σR2 | K264 | K371 | σR3-σR4 | K493 | Y571 | σR4 | D546 | K593 |
| σNER-σR2 | K264 | K418 | σR3-σR4 | K496 | K593 | σR3-σR4 | K462 | D613 |
| σNER-σR2 | K236 | K371 | σR3-σR4 | K493 | K593 | σR4 | D546 | K557 |
| σNER-σR2 | K343 | K371 | σR4 | K578 | K593 | σR4 | D546 | K593 |
| σNER-σR2 | K220 | K371 | σR4 | Y571 | K593 | | | |
| σNER-σR2 | K264 | K377 | σR4 | K593 | K597 | | | |
| σNER-σR3 | K220 | K462 | | | | | | |
| σNER-σR3 | K257 | K462 | | | | | | |
| σNER-σR3 | K264 | K462 | | | | | | |
| σNER-σR3 | K220 | K493 | | | | | | |
| σNER-σR3 | K343 | K462 | | | | | | |
| σNER-σR3 | K236 | K462 | | | | | | |
| σNER-σR3 | S241 | K493 | | | | | | |
| σNER-σR3 | K264 | K496 | | | | | | |

**Fig. 2.**
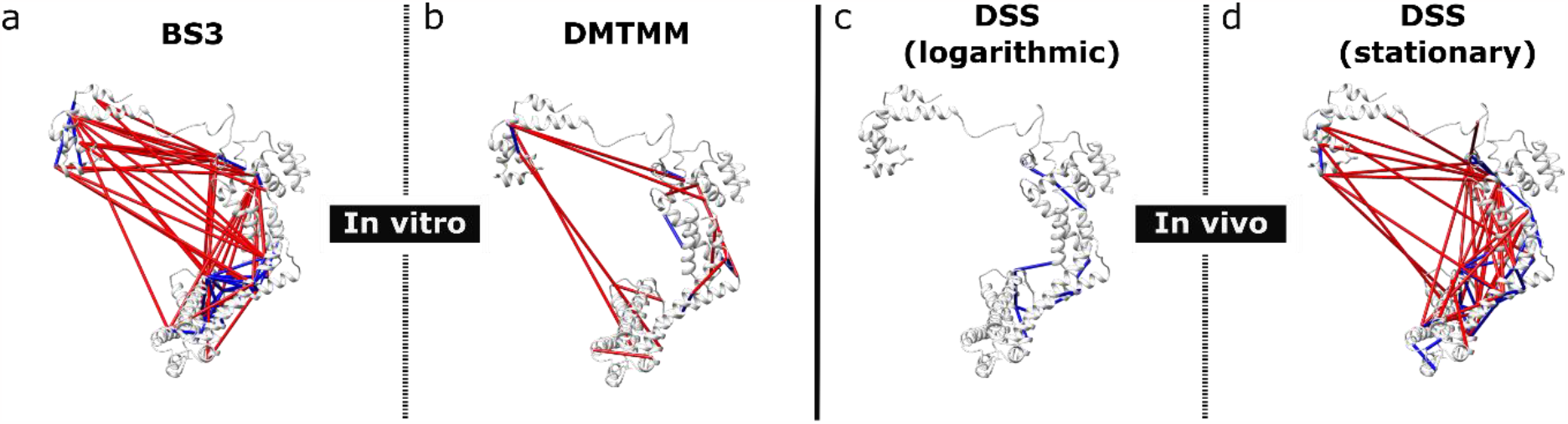
Apo-σ^70^ *in vitro* and *in vivo* distances recovered from CL-MS suggest a compact conformation exists in solution and during stationary phase of bacterial growth. The RNAP-bound σ^70^ conformation from PDB structure (6P1K). (**a**) *In vitro* BS^3^ and (**b**) DMTMM cross-links. (**c)** *In vivo* DSS cross-links at logarithmic growth phase. (**d**) *In vivo* DSS cross-links at stationary growth phase. Blue and red – cross-links recovered distances that are within or are not within the C_α_-C_α_ distance covered by the crosslinker, respectively.

**Fig. 3.**
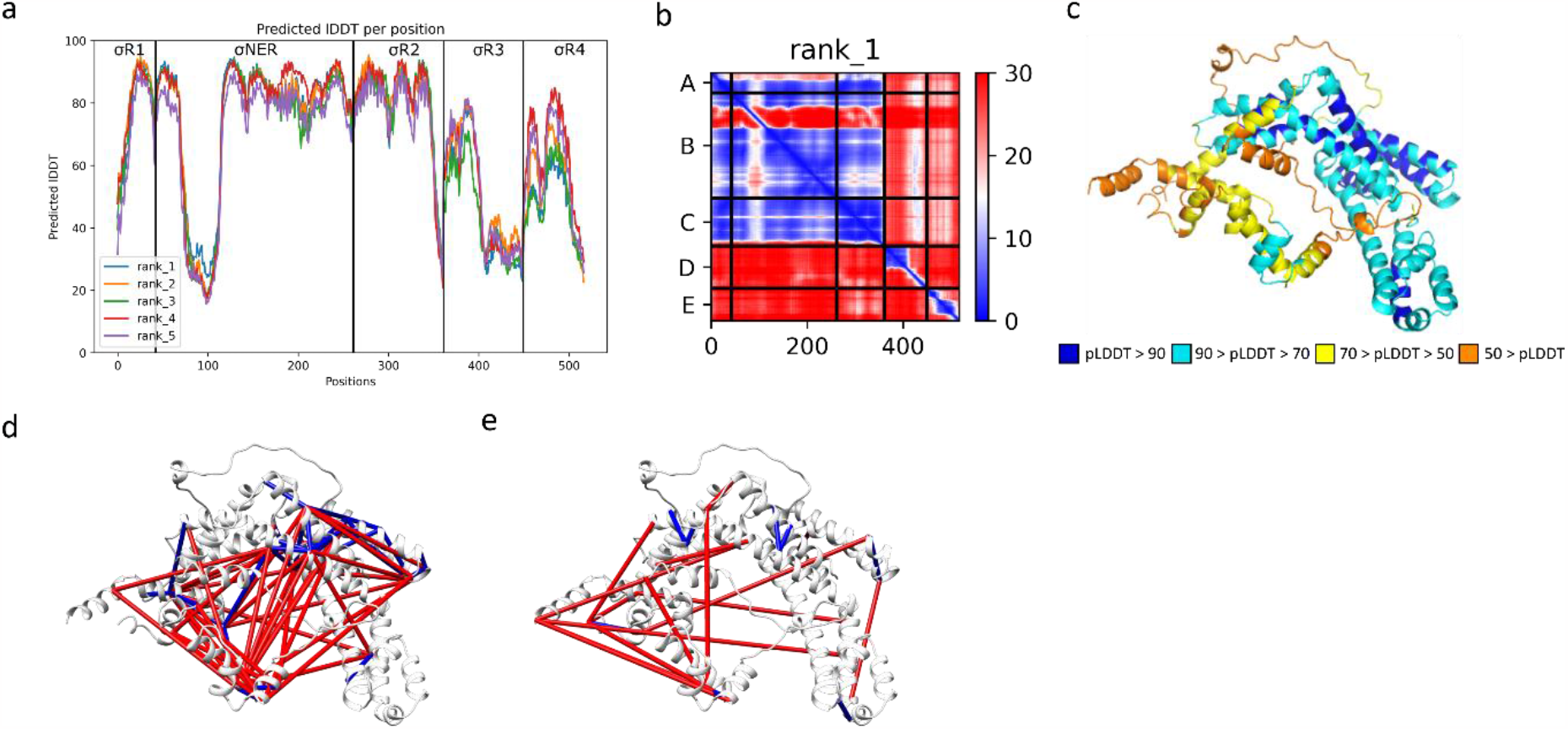
AlphaFold2 Multimer predicts a compact conformation which does not satisfy CL-MS data. (**a**) Predicted pLDDT (AlphaFold2 output) for each residue in the top 5 predicted model structures. While the domains are predicted with high LDDT, linkers are significantly lower in their values. σ^70^ domains are separated in different boxes. (**b**) Predicted inter-residue interactions of the top-ranking model, where A, B, C, D and E correspond to σR1, σNER, σR2, σR3 and σR4, respectively. Blue color indicates that interaction occurred between the two regions, while red color indicates no interaction. (**c**) Rank 1 relaxed model is in a more compact conformation compared to RNAP-bound σ^70^. (**d**) BS^3^ and (**e**) DMTMM cross-links displayed on the rank 1 relaxed model. Blue and red – cross-links recovered distances that are within or are not within the C_α_-C_α_ distance covered by the crosslinker, respectively.

### *In vivo* CL-MS identifies structural features of the compact apo-σ^70^ conformation

To explore whether unique features of the predominant compact conformation of apo-σ^70^ exist in living *E. coli* cells, we perform *in vivo* CL-MS during the logarithmic and stationary phases of bacterial growth. Disuccinimidyl suberate (DSS) is a crosslinker, which shares the same end-to-end length and cross-linking chemistry as BS^3^. Yet, unlike BS^3^, DSS can passively penetrate through cell membranes, including *E. coli*^63^. During the logarithmic phase, σ^70^ is expected to be found mostly in the RNAP-bound form, due to its engagement in the transcription of house-keeping genes^64^. Conversely, during the stationary phase, σ^38^ replaces σ^70^ in the transcription process, which suggest that most of σ^70^ is in a RNAP-unbound form, potentially in the apo form^65^. Following FDR-based filtration of the *in vivo* CL-MS results (Fig. S3, c), we show that during the logarithmic phase of bacterial growth, cross-links can be explained by the RNAP-bound σ^70^ extended conformation (Fig. 2, c). On the other hand, during the stationary phase, we capture DSS cross-links, which we previously captured *in vitro* for apo-σ^70^ using BS^3^, and are found to describe exclusively its predominant compact conformation (Fig. 2, a & d). Out of the 49 *in vitro* cross-links, which we found to be unexplained by PDB structures (Fig. 4, c-d, and Table S2), we recover 11 cross-links *in vivo* using DSS during the stationary phase (Fig. 4, b). The results of western-blot of enriched σ^70^ after *in vivo* DSS cross-linking indicate the cross-linked construct is mostly a monomeric σ^70^ with a small fraction of complexes or oligomers that include σ^70^ as well (Fig. S7). This, in turn, suggests that the majority of the recovered cross-links are a result of conformational changes in the monomeric apo-σ^70^. The cross-links recovered from *in vivo* CL-MS are summarized in supplementary spreadsheet S1 and are displayed on top of a representative PDB structure of σ^70^ (6P1K; Fig. 4). Our results suggest that this apo-σ^70^ compact structure in *E. coli* can be anticipated not during the logarithmic phase, where housekeeping genes are needed, but rather during stress when the binding of σ^70^ to RNAP is replaced by an alternative σ factor. Furthermore, we analyze the interaction interface of σ^70^ with DNA in the RP_o_ complex using the PDB structures summarized in Table S2 and the program PISA^66^. We find that part of σR1.2, σR2, σR3 and σR4’s DNA interacting residues in the RP_o_ stage undergo cross-linking in the apo-state using DMTMM, these residues include D96, K392, K493, K593, S389, K462. As mentioned DMTMM is a carboxy to amine coupler active at a distance of up to 16Å (C_α_-C_α_), hence in apo-state σ^70^ DNA binding residues are less accessible to the solvent and are less likely to form interaction with DNA. Based on our knowledge on the interaction of the transcription complex with promoter DNA, together with the fact that all structures of RNAP-bound full-length σ^70^ documented in the PDB are in the extended conformation, we suggest that the unique compact conformation of apo-σ^70^ might be responsible for the inhibition of DNA-interaction, and hence serve as a conformational switch activated upon binding to RNAP.

## Discussion

In this work, we provide evidence that apo-σ^70^ is found predominantly in a compact conformation, in which σR4 is in close proximity to σR2 and σR3. We perform CL-MS and further confirm the compact conformation of apo-σ^70^ both *in vitro* and *in vivo*, under stationary phase of the bacterial growth. Yet, an atomic resolution experimental structure of apo-σ^70^ has not yet been elucidated. CL-MS is a high throughput technique able to recover distance restraints between pairs of residues, from just a few micrograms of protein^48–53^.

CL-MS is limited in that it reports solely on pairs of residues that are within a given distance range and not on the pairs of residues that are at longer distances. Yet, we use cross-linkers to cover a variety of interresidue distances and to increase the accuracy of the suggested compact conformation and attain both short and intermediate range distances. The fragmentation score suggests how many times a certain peptide was detected. Above a threshold fragmentation score of 0.7 the FDR decreases dramatically. Nevertheless, as we show in the western-blot and in SEC-MALS measurements, our filtered cross-link list includes mostly ones that originate from the monomeric apo-σ^70^, while some cross-links probably originate from high-order oligomers or complexes at lower frequency. Overall, CL-MS results suggest that apo-σ^70^ may attain multiple different compact structural organizations, which are yet to be structurally recorded in the PDB.

Using smFRET, we show that in the presence of relatively high concentrations of promoter dsDNA, apo-σ^70^ remains predominantly in a compact conformation. Interestingly, a small fraction of σ^70^ in solution exhibits a conformation that is slightly less compact, suggesting that DNA binding may occur to some degree and might induce a slight conformational change between σR2 and σR4 towards a slightly open conformation. Therefore, it raises the question of whether or not the extended conformation of σ^70^ exists intrinsically and dynamically in apo form, interchanging between the two conformations. In fact, previous smFRET studies suggested that such intrinsic conformational dynamics may exist in *E. coli* apo-σ^70^ <u>^5^</u>. As previously shown by *Vishwakarma et al*.^37^, we measure the distance between σ^70^ regions using FRET efficiencies between a donor dye labeling residue 579 of σR4 and an acceptor dye labeling residue 442 of σR2. Overall, the results point towards a preexisting conformational equilibrium between a predominant compact conformation, and a minorly-populated less compact conformation of apo-σ^70^. Furthermore, we show that σ^70^ may partially bind dsDNA in the absence of RNAP, and at low affinity (μM), as opposed to DNA binding in the presence of RNAP, which occurs at high affinity (nM). In that respect, it is important to mention that σ^70^ has been reported to bind DNA structures that deviate from the dsDNA one (e.g., ssDNA in aptamers)^45^, however not to dsDNA, and not at high affinities. The ability of apo-σ^70^ to bind DNA is predictable since σ^70^ undergoes intrinsic dynamic conformational changes, which slightly open the σR2 and σR4 DNA binding interface. One can speculate that if σ^70^ would bind dsDNA not in the context of RNAP, it would probably bind it nonspecifically, hence the interaction would be unstable and will lead to inefficient transcription initiation. Indeed, our results encourage the hypothesis and show that dsDNA-induced σR2-σR4 conformational changes are small, occur only to a small fraction of σ^70^ molecules, and only at relatively high dsDNA concentrations. Therefore, to regulate the transcription process and allow efficient transcription initiation it is necessary to protect the DNA binding residues of σ^70^ from being exposed, until binding to RNAP occurs. Binding will activate the required conformational change in σ^70^, which will extend it and expose its DNA-binding residues. Moreover, similar to core RNAP, anti-σ factors bind σ^70^ covering the σR2-σR4 DNA-binding interface^1^. Therefore, the σR2-σR4 interface can serve as a conformational switch that self-inhibits and sterically protects the inner DNA binding residues from being exposed, until needed.

Next, we captured cross-links *in vivo* using DSS and compared it to the BS^3^ cross-links *in vitro*, since they both share similar thresholds. We found similarity in the DSS and BS^3^ crosslinks, which were found to be exclusive to the predominant compact conformation of apo-σ^70^, suggesting that the compact conformation exists also *in vivo*. Importantly, this finding reports on a biologically-relevant event, where apo-σ^70^ exists in living *E. coli* cells at stationary phase. In other words, σ^70^ is found in an unbound state and is not degraded, or at least not in full. Nevertheless, these results were achieved using a high copy number plasmid of σ^70^, and hence might also be influenced by over-expression effects. Yet, if over-expression would pose a problem, we would expect to observe the unique cross-linking signatures of the compact apo-σ^70^ conformation also in the logarithmic phase since at this stage σ^70^ exist in the cell at high concentration. To confirm that most cross-links arise from conformational changes, we have also shown that the majority of *in vivo* cross-linked purified σ^70^ (i.e., a fraction of 93%) is present in the monomeric form.

In summary, we propose that apo-σ^70^ organizes in a predominant compact conformation, which may lead to self-inhibition of the high-affinity promoter binding capabilities of σ^70^. Only upon activation by binding to RNAP, a large conformational change is anticipated, which exposes these DNA-binding residues. We provide biological evidence for the existence of the compact apo-σ^70^ conformation in live *E. coli* cells during the stationary phase.

## Acknowledgments

The plasmid of the Cys-less σ^70^ from which we prepared the doubly-labeled σ^70^ variant for our smFRET measurements, was provided to us as a gift from the laboratory of Dr. Shimon Weiss, UCLA. We would like to thank Dr. Paul David Harris for consultation regarding mpH^2^MM analyses of smFRET data. We would like to thank Dr. Shimon Weiss for fruitful discussions and for inspecting the text of this work. Molecular graphics and analyses performed with UCSF ChimeraX, developed by the Resource for Biocomputing, Visualization, and Informatics at the University of California, San Francisco, with support from National Institutes of Health R01-GM129325 and the Office of Cyber Infrastructure and Computational Biology, National Institute of Allergy and Infectious Diseases.

## Author contributions

Analytical tools: NK

Protein purification: KJ

Experimental data: KJ, JZ

Analysis of experimental data: KJ, EL

Writing – original draft: EL, KJ

Writing – reviewing & editing: NK, JZ

## Data and materials availability

SmFRET data – https://doi.org/10.5281/zenodo.7173886^67^. CLMS data – available at Proteomics Identification Database (PRIDE), The structural basis for the self-inhibition of DNA binding by apo-σ^70^ - CLMS data (ebi.ac.uk; PXD037183).

