## Supplementary Information for "Evidence for a compact σ^70^ conformation *in vitro* and *in vivo*"

**This PDF file includes:**

Materials and Methods

Figures S1 to S5

Tables S1 & S2

### Materials and Methods

#### *Recombinant protein expression and purification*

Plasmid encoding for the RNAP holoenzyme rpoD ($\sigma^{70}$) subunit was obtained from Addgene as a gift from Dr. Irina Artsimovitch (Addgene plasmid #104399)[68]. Plasmid pET28b(+)_reverse_comp_rpoD(-C) expressing the *E. coli* $\sigma^{70}$ without cysteines was a gift from the laboratory of Shimon Weiss at UCLA. Purification is performed following previously published protocol[27,68] with minor adjustments. Briefly, plasmids are transformed into *E.coli* BL-21 competent cells. A single colony of the transformed BL-21 cells is suspended in 10 mL of LB media containing 50 µg/mL of Kanamycin and incubated overnight at 37 °C and 250 rpm. Overnight culture is then diluted (1:100) into a newly autoclaved 2 L erlenmeyer flask, containing 1 L of LB media and supplemented with 50 µg/mL of Kanamycin. Culture growth is monitored by periodically measuring $OD_{\lambda=600nm}$; once $OD_{\lambda=600nm}$ reached 0.6 (after ~4 hours), recombinant protein expression is induced with IPTG addition at a final concentration of 1 mM. After 3 hours of incubation at 37 °C and 250 rpm, cells are harvested by centrifuging at 6,000 g and 4 °C . The bacterial pellet is resuspended on ice in lysis buffer (50 mM HEPES-KOH (pH 7), 500 mM NaCl, 5% glycerol) containing 1 tablet of cOmplete™ EDTA-free protease inhibitors cocktail (Roche). Cell suspension is then supplemented with lysozyme to a final concentration of 1 mg/mL. Cells are then disrupted by ultrasonication on ice at 60% amplitude for 10-12 cycles (20 second pulses with 50 second intervals). Cell debris was removed by centrifuging at 12,000 rpm and 4 °C for 30 minutes. Then for an additional 15 minutes after removing the pellet. Ni-Sepharose 4 mL column is used to separate the His-tagged $\sigma^{70}$ from the supernatant, with all washings and elution performed at 4 °C. Fractions suspected of containing $\sigma^{70}$ are collected and run in SDS-PAGE for purity assessment. Pure fraction containing high concentration of $\sigma^{70}$, are combined and further purified by running through Resource 15Q 1 mL column for ion exchange chromatography. Pure fractions are assessed by running SDS-PAGE and Coomassie staining. Then, pure fractions are dialyzed against dialysis buffer containing (20 mM HEPES-KOH (pH 7.5), 200 mM NaCl, 0.2 mM DTT and protease inhibitor cocktail). Protein concentration is then determined and protein is stored in 50% glycerol at 20 °C.

#### *$\sigma^{70}$ mutagenesis and double labeling*

Point mutations are introduced to the cys-less *E. coli* $\sigma^{70}$ plasmid, pET28b(+)_reverse_comp_rpoD(-C), at positions 442 and 579 following a 2-step PCR protocol[68] and using a set of specific primers. Mutagenesis results are confirmed by Sanger sequencing and comparison. Purification of the mutant $\sigma^{70}$ containing two cysteines at positions 442 and 579, is performed similarly to the *wt* $\sigma^{70}$ purification described above. For labeling, mutant $\sigma^{70}$ is dialyzed against (50 mM HEPES-KOH pH=7.2 and 0.5 mM EDTA) overnight at 4 °C to reduce the glycerol concentration. After dialysis, a final concentration of 1.25 µM mutant or *wt* $\sigma^{70}$ are suspended in (50 mM HEPES-KOH pH=7.2, and 0.1 mM EDTA) at final volume of 1 mL with 5 µM of Tris(2-carboxyethyl)phosphine (TCEP), to reduce any disulfide bridges without introducing free reacting thiols, for a more efficient labeling. The reaction is stirred for 45 minutes at room temperature. ATTO 643 and ATTO 550, with linkers to an iodoacetamide group, are then added to a final concentration of 1.25 µM, in a dark room and the reaction is stirred at room temperature for 4-5 hours. The reaction is then transferred to 4 $^{o}$C overnight. To terminate the reaction, β-mercaptoethanol is added to a final concentration of 5 µM at ~1 mL final volume and incubated with stirring for 30 minutes at room temperature. The sample is then dialyzed against (50 mM HEPES-KOH (pH=7.2), and 0.1 mM EDTA) overnight at 4 $^{o}$C, and stored at -20 $^{o}$C with 50% glycerol.

***RNAP holoenzyme transcription assay***

While in the absence of an RNA transcript the ssDNA FRET probe show a mean FRET efficiency of ~0.5 (Fig. S1, b), once a transcript is introduced the ssDNA FRET probe undergoes hybridization, which stretches the DNA probe, leading to a decrease of the FRET efficiency to ~0.0 (Fig. S1, d). Therefore, the decrease in the FRET efficiency indirectly indicates the presence of transcripts that were successfully synthesized, hence active transcription complex and proper binding of $\sigma^{70}$ to both RNAP and promoter.

***SmFRET measurements and data analysis***

SmFRET measurements are performed on dually labeled mutant $\sigma^{70}$ on a confocal-based microscopy setup (ISS$^{TM}$ , Champaign, IL, USA) assembled on top of an Olympus IX73 inverted microscope stand (Olympus, Tokyo, Japan). Samples are measured in a high glass bottom µ-slides (Ibidi), with an acquisition time of 1 hour per technical repeat at room temperature (23 $^{o}$C.). We use 532±1 nm (FL-532-PICO, CNI, China) and a 640±1 nm (QuixX ® 642-140 PS, Omicron, GmbH) pulsed picosecond fiber and diode lasers, respectively, operating at 20 MHz repetition rate. Scattering and fluorescence photons return in the excitation path, collected through the same objective. Then, while scattering photons are reflected, fluorescence photons are transmitted through a major dichroic mirror with high reflectivity at 532 nm and 640 nm (ZT532/640rpc, Chroma, Bellows Falls, Vermont, USA) and is focused with an achromatic lens (25 mm Diameter x 100 mm FL, VIS-NIR Coated, Edmund Optics) onto a 100 µm diameter pinhole (variable pinhole, motorized, tunable from 20 µm to 1 mm, custom made by ISS$^{TM}$), and then re-collimated with another achromatic lens (AC254-060-A, Thorlabs). Fluorescence originating from either donor or acceptor dyes is then split into two detection channels using a 605 nm cutoff dichroic mirror, and is then further cleaned using a 698/70 nm bandpass filter for acceptor emission and a 585/40 nm bandpass filter for donor emission. Single fluorescence photons are then detected using hybrid PMTs (Model R10467U-40, Hamamatsu, Japan), and single photon detection event time tags are collected using a time-correlated single-photon counting card (SPC 150N, Becker & Hickl, GmbH). All measurements are performed in the same buffer (20 mM HEPES-KOH at pH 7.0, 50 mM KCl, 10 mM $MgCl_2$, 2 mM β-mercaptoethanol). SmFRET experiments are performed using nanosecond alternating laser excitation, nsALEX[69], also known as pulsed interleaved excitation, PIE[70]), which provides information on the excitation origin of the donor and acceptor photons. Data is then analyzed using FRETbursts[71] and mpH$^2$MM[56]. A burst identified using a dual channel burst search[72] is considered a single molecule event, only if for each consecutive 10 photons a count rate of at least 16 times higher than the background rate, and only if it includes at least 30 photons originating from donor excitation and at least 30 photons originating from acceptor excitation.

#### *In vitro single-molecule FRET-based transcription activity of the purified recombinant RNAP*

The smFRET based transcription activity assay is performed as previously mentioned[54,55]. Briefly, The RNA polymerase is incubated in KG7 buffer (40 mM HEPES (KOH), pH=7.0, 10 mM $MgCl_2$, 1 mM DTT, 5% glycerol, supplemented with 1 mM TROLOX and 10 mM MEA) with the mutant-$\sigma^{70}$ (442C, 579C) at 37 $^oC$ for 30 minutes to form the holoenzyme complex. Next, linear dsDNA *lacCONS* promoter, which promotes the synthesis of a nascent RNA transcript with 20 adenine (20A; Fig. S1a) bases at the 3'-end, is introduced to the holoenzyme complex at 37 $^oC$ to form the RNAP-promoter open complex. Later, all nucleotides are introduced together with RNase inhibitor (BioLabs), and the sample is again incubated at 37 $^oC$. guanidinium-HCl is then added at room temperature (25 $^oC$ ) to quench the reaction. Then, a ssDNA probe consisting of 20 deoxy-thymine (20dT) and labeled by a donor dye (ATTO 488) at the 5'-end, and an acceptor dye (ATTO 647N) at the 3'-end, is used for detecting nascent RNA transcripts via hybridizing to them, inducing a reduction in the end-to-end FRET values relative to that in the free ssDNA FRET probe. In this manner, the probe is able to produce FRET signals as long as it is not degraded. If the probe is degraded the FRET signal efficiency would be reduced to E=0. A final concentration of 50 pM of the probe is incubated in 100 µL of the prepared sample. If RNA transcripts are produced, we expect to detect a decrease in the FRET efficiency due to hybridization of the 20A bases of the nascent RNA transcript to the 20dT bases of the probe.

#### *In vitro Cross-linking and mass-spectrometry preparation*

*In vitro* CL-MS is performed as previously described[48,73]. Briefly, $BS^3$ powder is used to prepare a 10 mM stock solution using HEPES buffer (20 mM HEPES (KOH) pH=7.0, 50 mM KCl, 10 mM $MgCl_2$ and 2 mM 2-mercaptoethanol). Similarly, DMTMM is prepared at a 70 mM stock solution. For cross-linking, a final concentration of 1 mM $BS^3$ or 7 mM DMTMM is incubated with 5-10 µg of *wt* $\sigma^{70}$ at 30 $^oC$ for 1.5 hours with shaking at 600 rpm. Next, to terminate the reaction, ammonium bicarbonate is added at 3X the concentration of the crosslinker from a 1 M stock solution (so the sample is not diluted), and incubated at 25 $^oC$ for 30 minutes with shaking. Later, preparation of the samples for mass spectrometry and mass spectrometry RAW data files analysis, is performed as previously described[73,74] with minor adjustments, see below the section on *crosslinks identification and estimation of* false detection rate (*FDR*).

#### *In vivo Cross-linking and mass-spectrometry preparation*

Plasmid encoding for the RNAP holoenzyme rpoD ($\sigma^{70}$) subunit was obtained from Addgene as a gift from Dr. Irina Artsimovitch (Addgene plasmid #104399)[68]. After transformation and isolation of a single colony containing the plasmid as described above (Recombinant protein expression and purification), an overnight culture is diluted (1:100) into a newly autoclaved 250 mL erlenmeyer flask, containing 100 mL of LB media and supplemented with 50 µg/mL of kanamycin. The flask is incubated at 37 $^{o}$C with shaking until $OD_{\lambda=600nm}$ reaches 0.6, then IPTG is added and the flask is incubated again at 37 $^{o}$C. After 3 hours a bacterial pellet from 5 mL of bacterial culture is collected and stored at -80 $^{o}$C. The flask is incubated again at 28 $^{o}$C with shaking. After overnight incubation, bacterial growth was closely monitored by measuring $OD_{\lambda=600nm}$ every 15 minutes. Once $OD_{\lambda=600nm}$ values stop elevating (stationary phase; monitored periodically using a spectrophotometer) a bacterial pellet from 5 mL of bacterial culture is collected and stored at -80 $^{o}$C. For the cross-linking reaction a 250 mM stock solution of DSS is prepared in DMSO. The collected bacterial pellets are resuspended in lysis buffer (50 mM HEPES (KOH) pH=7.0, 500 mM NaCl and 5% glycerol) containing 10 mM DSS to increase membrane permeability. First, DSS is resuspended from the stock solution in 1 mL of lysis buffer, since DSS is not completely soluble in water it forms a precipitate, and we take only the dissolved fraction which is ~1 mL of lysis buffer supplemented with a final concentration of ~10 mM DSS. The cross-linking reaction is then incubated at 30 $^{o}$C for 20 minutes with shaking. Later, the cross-linking reaction is quenched by adding 40 mM final concentration of ammonium bicarbonate and incubating at room temperature (~25 $^{o}$C ) with shaking for another 20 minutes. Centrifuge at 6,000 g and 4 $^{o}$C for 5 minutes to collect the cells, pellets are then stored at -80 $^{o}$C. Pellets are resuspended in 1 mL of lysis buffer, and lysed by sonication using the Qsonica 422 Ultrasonic probe at 60% Amp for 12 cycles of 10 seconds ON and 25 seconds intervals. Centrifuge to discard cell debris, keep the supernatant. Ni-Sepharose high performance beads (GE healthcare) are washed thoroughly with lysis buffer to rinse off the ethanol. The supernatant is then incubated with 10 µL of the Ni beads and incubated at 4 $^{o}$C with minimal shaking (to prevent sinking of the beads) for 4 hours. Following, samples are centrifuged to take out the flow-through, and the beads are resuspended with wash buffer (lysis buffer + 20 mM Imidazole), centrifuge at 950 g and 4 $^{o}$C for 2 minutes and take out the liquid, the washing process is repeated three times. For elution, the beads are then resuspended in 50 µL of elution buffer (lysis buffer with 400 mM Imidazole), and kept at room temperature for 30 minutes while inverting the tubes every 2 minutes to prevent beads from sinking to the bottom. Centrifuge at 950 g and 4 $^{o}$C for 2 minutes, and collect the liquid without disrupting the beads. Then samples are prepared for mass spectroscopy as previously described[73,74]. Briefly, the protein is precipitated in 1 mL of acetone (-80 $^{o}$C ) for 1 hour, followed by centrifugation at 14,000 g. The pellet is resuspended in 20 mL of 8 M urea. The urea is diluted by adding 200 mL of digestion buffer (25 mM TRIS, pH = 8.0; 10% Acetonitrile). We add 0.5 mg of trypsin (Promega, Madison, Wisconsin) to the diluted urea and digest the protein overnight at 37 $^{o}$C under agitation. Following digestion, the peptides are desalted on C18 stage-tips and eluted by 55% acetonitrile. The eluted peptides are dried in a SpeedVac, reconstituted in 0.1% formic acid, and measured in the mass spectrometer.

#### *Mass spectrometry analysis*

The samples are analyzed by a 120-min 0-to-40% acetonitrile gradient on a liquid chromatography system coupled to a Q-Exactive HF mass-spectrometer. The analytical column is an EasySpray 25 cm heated to 40 $^{o}$C. The method parameters of the run are as follows: Data-Dependent Ac-quisition; Full MS resolution 70,000 ; MS1 AGC target 1e6; MS1 Maximum IT200 ms; Scan range 450 to 1,800; dd-MS/MS resolution 35,000; MS/MS AGCtarget 2e5; MS2 Maximum IT 600 ms; Loop count Top 12; Isolation window1.1; Fixed first mass 130; MS2 Minimum AGC target 800; Peptide match - off; Exclude isotope - on; Dynamic exclusion 45 seconds. Each cross-linked sample is measured twice in two different HCD energies (NCE): 26, and stepped 25, 30, and 35. All cross-linked samples are measured with the following charge exclusion: unassigned,1,2,3,8,>8. Proteomics samples are measured with the following charge exclusion: unassigned,1,8,>8.

#### *Cross-links identification and filtration*

The RAW data files are converted to MGF using Proteome Discoverer (Thermo). Then, FindXL[48] is used to exhaustively enumerate all the possible peptide pairs originating from $BS^3$ or DMTMM crosslinks, with the following search parameters: i) Sequence database - $\sigma^{70}$ sequence taken from UniProt (P00579); ii) Protease – trypsin, allowing up to three mis-cleavage sites; iii) Variable modifications: methionine oxidation, lysine with hydrolyzed mono-link; iv) Cross-linking must occur between two lysine residues or lysine and glutamic acid/aspartic acid for ($BS^3$ /DSS or DMTMM, respectively); v) Cross-linker is not cleaved; vi) MS/MS fragments to consider: b-ions, y-ions; vii) MS1 tolerance – 6 ppm; viii) MS2 tolerance – 8 ppm; and ix) Cross-linker mass – one of three possible masses: 138.0681, 138.0681 + 1.00335, and 138.0681 + 2.0067 for the $BS^3$ /DSS and -18 Da for DMTMM which represent the release of a water molecule. The three masses address the occasional incorrect assignment of the mono-isotopic mass by the mass spectrometer.

#### *Estimation of FDR*

The FDR is estimated by repeating the cross-link identification analysis 20 times with an erroneous cross-linker mass of 138.0681 × N/138 Da (for $BS^3$ and DSS), where N = 160, 161, 162, ... 179. This leads to bogus identifications with fragmentation scores that are generally much lower than the scores obtained with the correct cross-linker mass. For the identification of true cross-links, we set the threshold on the fragmentation score according to the desired FDR value (Fig. S5). For DMTMM, same as mentioned above with the erroneous cross-linker mass of -18.01056 × N/18 Da (Figs. S5-S7). With the fragmentation scores 1 and 2 set to 2.0 and 0.7, respectively, we get a FDR of 0.65% for BS3 (Fig. S5), 1.6% for DMTMM (Fig. S6) and 0.55% for DSS (Fig. S7) cross-links.

#### *Structure modeling*

Using AlphaFold, RosettaFold and OmegaFold we submitted a single sequence of the entire $\sigma^{70}$, and got an output of different structure models ranked according to the scoring parameters of each algorithm. However, using ColabFold version 1.5.2, a protein complex structure prediction using AlphaFold2-multimer version 3 (v3), we submitted each region of the $\sigma^{70}$ as a separate sequence, allowing AlphaFold2-multimer v3 to reorganize the structure of apo-$\sigma^{70}$ from rigid domains taking into account the high flexibility of the linkers. Using MMseqs2 sequence alignments were generated, which created the data library that Colabfold used to predict the structure model. We used the "unpaired_paired" pair mode to pair sequences from the same species but also from different species as well. This increases the model prediction accuracy, and is possible due to the high conservation rate between housekeeping σ factors of different species. We performed three recycles of the run to increase the validity of the results.

***Western-blot***

*In vivo* cross-linking and lysis were performed as described above. The lysate was incubated with Ni-beads for 4 hours at 4 $^{o}$C. Next the solution was centrifuged at 800 g to precipitate the Ni-beads binding $\sigma^{70}$, and the supernatant was separated from the precipitate. Then the beads were washed three times with wash buffer to get rid of all nonspecific Ni binders. Finally, the $\sigma^{70}$ was eluted using elution buffer. The elution was run in a 10% acrylamide SDS-PAGE and transferred to a PVDF membrane. Detection of $\sigma^{70}$ was performed by incubating the membrane at 4 $^{o}$C overnight in a 0.5 µg primary antibody solution, mouse IgG2b from Bio-legend, followed by an alkaline phosphatase tagged secondary antibody.

### Supplementary figures and tables

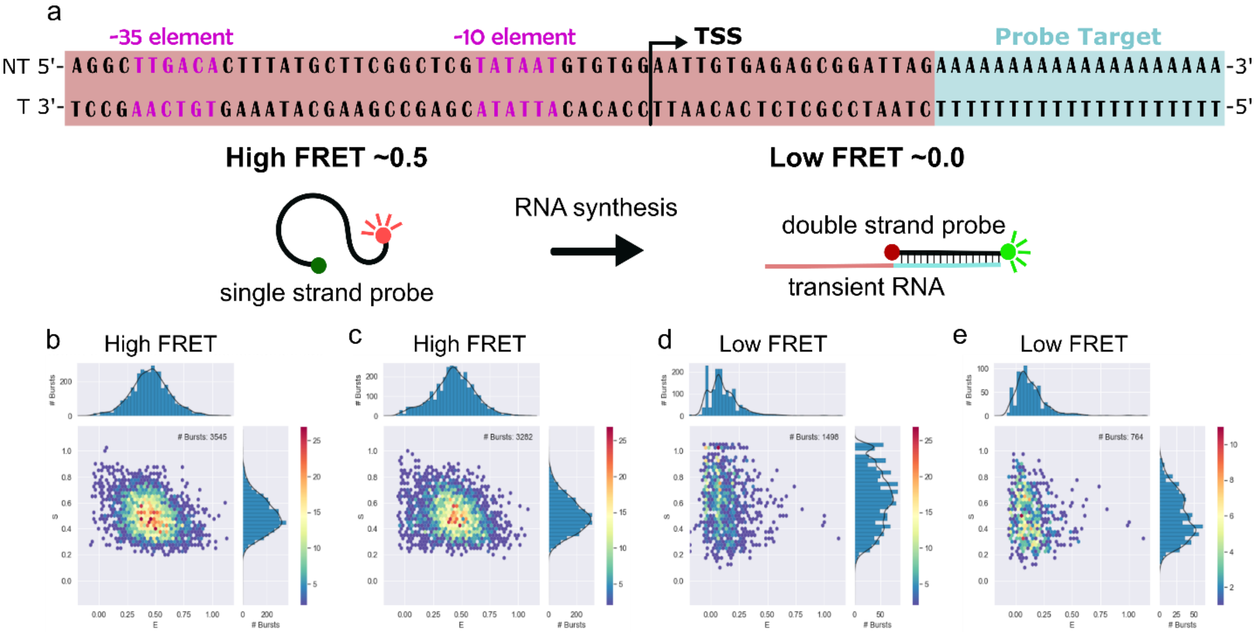


**Fig. S1. The RNAP holoenzyme of *wt* and mutant-σ$^{70}$ is transcriptionally active.** (**a**) LacCONS promoter sequence including a probe target sequence, and illustration of the 20dT FRET probe labeled at 3'- and 5'-ends with ATTO 488 and ATTO 647N, respectively. When the probe binds transient RNA, the binding fully stretches the probe. (**b**) 25 pM of the 20dT probe. A single mid-FRET sub-population exists, describing the short distance between the edges of the DNA probe. (**c**) 25 pM of the probe mixed with 1 nM of dsDNA lacCONS promoter, which should not induce a single-stranded DNA or RNA that hybridizes to the 20dT DNA probe, and hence the mid-FRET sub-population is similar to the one in panel a. 25 pM of the probe mixed with 1 nM of dsDNA lacCONS promoter, all four NTPs and RNAP holoenzyme of the mutant-σ$^{70}$ (**d**) and *wt* σ$^{70}$ (**e**). At this condition, a low-FRET sub-population exists, describing the long distance between the now stretched edges of the DNA FRET probe, upon hybridization with the transcribed RNA.

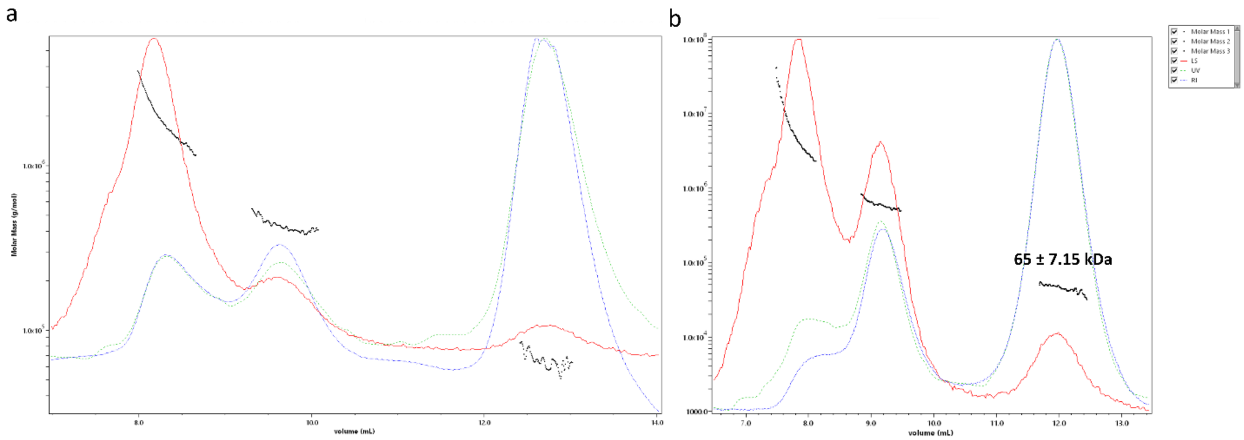


**Fig. S2. Size-exclusion chromatography of cross-linked and non-cross-linked samples of σ$^{70}$ show that σ$^{70}$ remains mostly in a monomeric conformation even after cross-linking.** SEC-MALS report of a non-cross-linked apo-σ$^{70}$ sample (**a**) and a cross-linked apo-σ$^{70}$ sample (**b**). Where the monomeric form elutes at ~12 mL and higher order oligomers elute between 6-10 mL depending on the size.

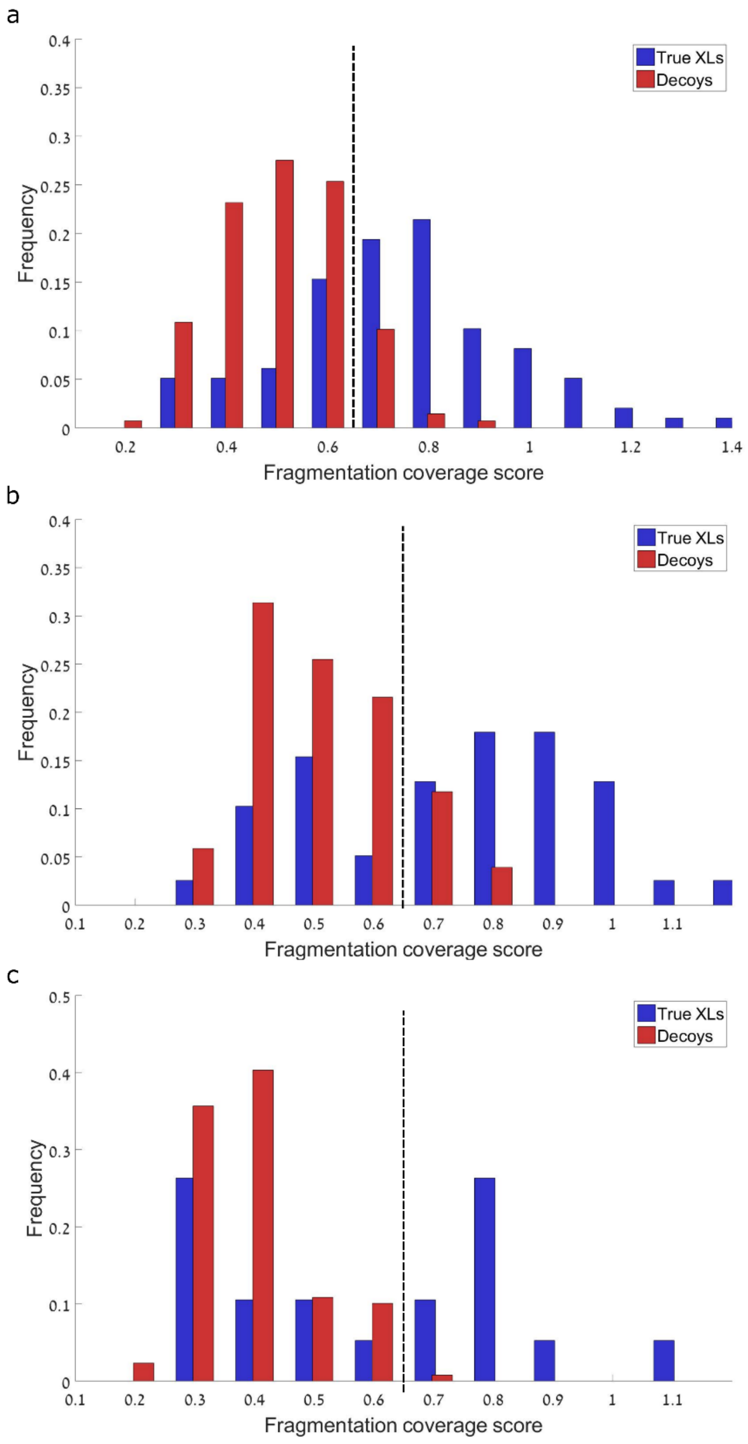


**Fig. S3. The frequency of true crosslinks vs decoy crosslinks as a function of coverage score.** $BS^3$ (**a**), DMTMM (**b**) and DSS (**c**). Above the fragmentation coverage score of 0.7 used in our analysis, the rate of decoy cross-links decreases drastically.

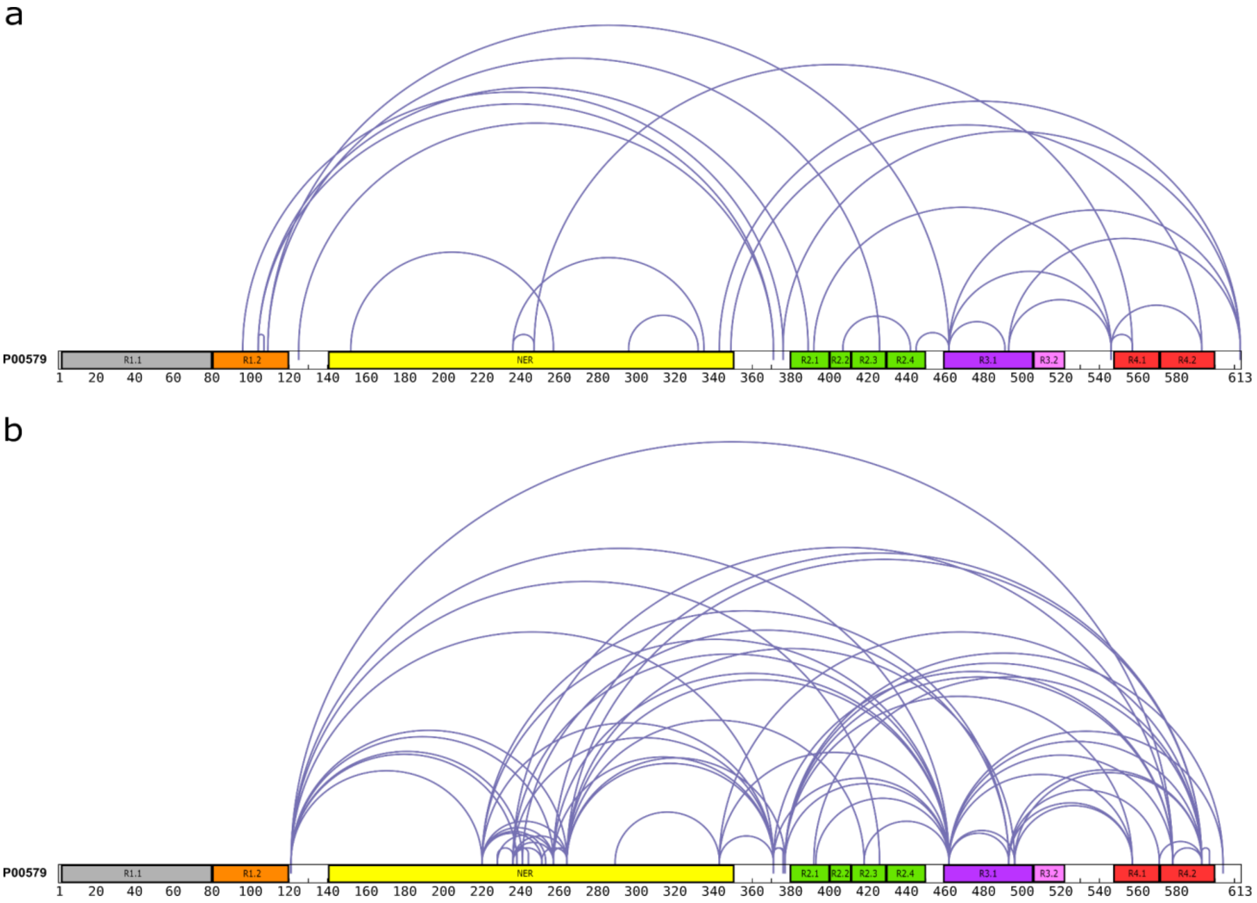


**Fig. S4. Arcs representation of *in vitro* crosslinked pairs of residues of apo-$\sigma^{70}$.** (**a**) $BS^3$ and (**b**) DMTMM cross-links**.** The $\sigma^{70}$ sequence was derived from UniProt (P00579). $\sigma^{70}$ regions are highlighted in color - σR1.1 in gray, σR1.2 in orange, σNER in yellow, σR2 in green, σR3.1 in magenta, σR3.2 in pink and σR4 in red.

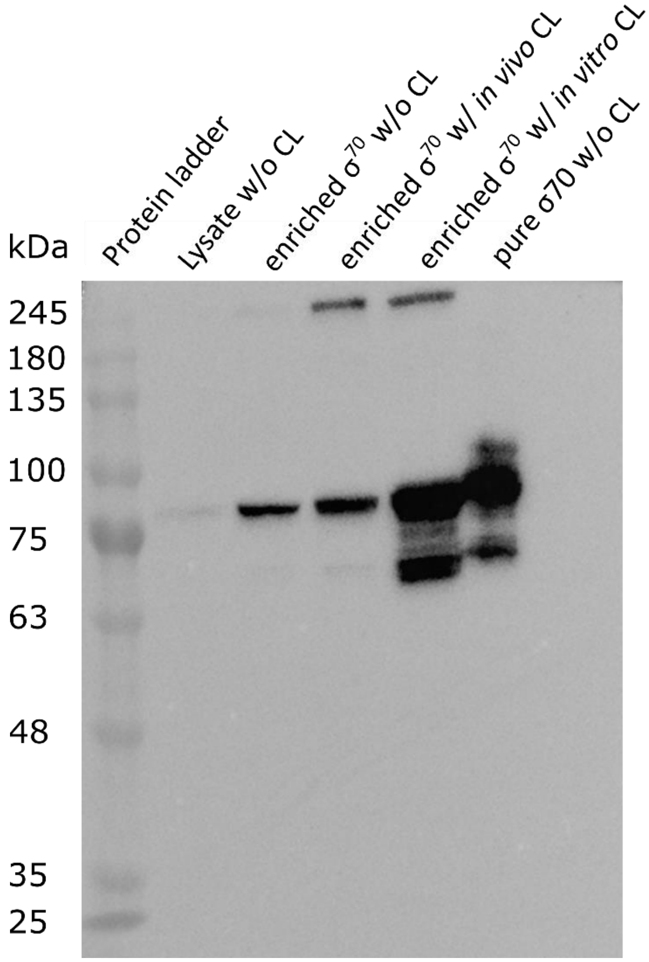


**Fig. S5. Western blot of *In vivo* and *In vitro* $\sigma^{70}$ samples.** Lanes are numbered from 1 to 6. Lane 1 is proteins ladder, lane 2 is lysate without cross-linking, lane 3 is enriched $\sigma^{70}$ without cross-linking, lane 4 is enriched $\sigma^{70}$ after cross-linking *in vivo*, lane 5 is enriched $\sigma^{70}$ cross-linked *in vitro* and lane 6 is purified $\sigma^{70}$ without cross-linking.

**Table S1. Transition rates of conformational dynamics revealed in the mpH$^2$MM analyses of *in vitro* smFRET measurements.** Transition times are given in ms, and the rates are calculated: $v = \frac{1}{transition\ time(ms)} * 1000$ , where v is the rate in s$^{-1}$. The table refers to FRET sub-populations referred to in Fig. 1 as blue-, green-, or red-colored. Bolded values are rate constant values for within-burst dynamics that is anything above 20 s$^{-1}$, due to bursts with durations shorter than 50 ms.

| **apo-σ$^{70}$** | | | |
|---|---|---|---|
| **From/To** | **Blue** | **Red** | **Green** |
| **Blue** | - | 9 | **42** |
| **Red** | 7 | - | **20** |
| **Green** | **35** | **22** | - |
| **σ$^{70}$ + 100 nM *lacCONS* promoter** | | | |
| **From/To** | **Blue** | **Red** | **Green** |
| **Blue** | - | 15 | **41** |
| **Red** | 18 | - | **320** |
| **Green** | **27** | **177** | - |
| **σ$^{70}$ + 2 µM *lacCONS* promoter** | | | |
| **From/To** | **Blue** | **Red** | **Green** |
| **Blue** | - | 18 | **23** |
| **Red** | **33** | - | **157** |
| **Green** | **20** | **69** | - |

**Table S2. Comparison of *in vitro* BS$^3$ and DMTMM apo-σ$^{70}$ cross-links against σ$^{70}$ structure models.**

| Conformation | PDB structure | Total BS$^3$ XLs | BS$^3$ XLs satisfaction (%) | Total DMTMM XLs | DMTMM XLs satisfaction (%) |
|---|---|---|---|---|---|
| Holoenzyme | 4JK1 | 68 | 44.1 | 20 | 30.0 |
| | 4JK2 | 68 | 44.1 | 20 | 30.0 |
| | 4LJZ | 58 | 43.1 | 19 | 36.8 |
| | 4LK0 | 58 | 43.1 | 19 | 36.8 |
| | 4LK1 | 58 | 43.1 | 19 | 36.8 |
| | 4MEY | 58 | 44.8 | 16 | 25.0 |
| | 4YG2 | 58 | 43.1 | 19 | 36.8 |
| | 6C9Y | 58 | 44.8 | 19 | 26.3 |
| | 6N57 | 58 | 43.1 | 19 | 36.8 |
| | 6N58 | 58 | 44.8 | 19 | 31.6 |
| | 6P1K | 58 | 43.1 | 19 | 31.6 |
| RP$_o$ | 6OUL | 58 | 44.8 | 19 | 36.8 |
| | 6PSQ | 58 | 43.1 | 19 | 36.8 |
| | 6PST | 58 | 43.1 | 19 | 36.8 |
| | 6PSS | 58 | 43.1 | 19 | 36.8 |
| | 6PST | 58 | 44.8 | 19 | 36.8 |
| | 6PSU | 58 | 44.8 | 19 | 31.6 |
| | 6PSV | 58 | 43.1 | 19 | 26.3 |
| | 6PSW | 58 | 44.8 | 19 | 36.8 |
| | 6WMU | 58 | 44.8 | 19 | 36.8 |
| | 7C97 | 58 | 43.1 | 19 | 36.8 |
| | 7CH2 | 58 | 43.1 | 19 | 36.8 |
| | 7DY6 | 58 | 43.1 | 19 | 36.8 |
| | 7MKD | 58 | 43.1 | 19 | 31.6 |
| | 7MKE | 58 | 43.1 | 19 | 31.6 |
| | 7MKI | 58 | 44.6 | 19 | 31.6 |
| | 7MKJ | 58 | 43.1 | 19 | 36.8 |
| Initially transcribed complex | 4YLN | 68 | 45.6 | 21 | 28.6 |
| | 4YLO | 68 | 45.6 | 21 | 38.1 |
| | 4YLP | 68 | 45.6 | 21 | 38.1 |
| RoseTTAFold | - | 68 | 44.1 | 25 | 28.0 |
| AlphaFold2 Multimer | - | 68 | 50.0 | 25 | 36.0 |
